# Thermophoresis of Molecules and Structures of different Sizes in Self-assembled Biomatrices

**DOI:** 10.1101/2022.04.11.487957

**Authors:** Ping Liu, Weilin Lin, Fabian Abele, Marcel Hanke, Yang Xin, Adrian Keller, Yixin Zhang

## Abstract

Upon subjecting biomolecules to non-equilibrium conditions, many biochemical and biophysical features such as biomolecular diffusion, protein folding, interaction kinetics, as well as enzyme-catalyzed reactions can be characterized in an aqueous solution. However, most assays under non-equilibrium conditions cannot be performed in complex self-assembled biomatrices (e.g. extracellular matrices) due to the limitations associated with sample handling, reaction design, and optical detection. Herein, we report the study of biomolecular thermodiffusion in non-covalently assembled synthetic or naturally derived hydrogels. This approach has been demonstrated with a large variety of analytes, including small molecules, polysaccharides, DNAs, DNA origami, and proteins in various polymer networks. The in-biomatrix method has also shown advantages over in-solution measurements: First, it allows us to analyze biomolecules in 3D matrices in a high-throughput fashion. Second, the aggregation of analytes can be remarkably prevented. Although the underlying physics of thermodiffusion is still not well-understood, we demonstrated that the thermodiffusion of surrounding networks will enhance the thermodiffusion of the analyte, an effect counteracting the hindered movement by the polymer network.

## Introduction

Thermophoresis (thermodiffusion or Soret effect), the phenomena of molecules drifting along temperature gradients, has been known for a long time^1,2^. Although the underlying physics remains poorly understood^3^, it has recently been explored as an all-optical technique to manipulate molecules as well as to measure molecular interactions in an aqueous solution^4–6^. A small temperature gradient of a few Kelvin (K) can be generated by IR laser irradiation at micrometer scales (approx. 100 *µ*m in diameter in this study) to induce thermophoresis, and the ensemble behavior of fluorescently labelled molecules can be followed. Moreover, microscale thermophoresis (MST) experiments can be carried out in a high-throughput fashion using commercialized instrumentation known as NanoTemper.

Like most biochemical and biophysical measurements, the MST assay in solution cannot fully represent the native environment of biomolecules such as in extracellular matrices (ECM). ECM can be viewed as self-organized and self-assembled polymer networks containing amphiphilic and zwitterionic biomacromolecular structures. The intracellular environments are also highly viscous or gel-like networks with different compartmentalization. In addition to compartmentalization by a lipid membrane, evidence is now mounting that liquid-liquid phase separation (LLPS) can lead to the formation of membrane-less compartments in cells^7^. The interactions between the signaling molecules and hydrated networks play crucial roles in the orchestrated activities of different cells. Ions, nutrients, growth factors, low-molecular-weight hormones, as well as therapeutic reagents diffuse through ECM and regulate the cell fates. Accordingly, the biophysical processes of the biomolecules have drawn the researchers’ attention. However, preparing a sample with the analyte homogeneously encapsulated in a hydrogel is far more complicated than preparing an aqueous solution. For example, many crosslinking chemistries are not compatible with proteins and cells. Moreover, to add reagents to a hydrogel, the additional solvent cannot be effectively mixed, thus preventing the creation of a controlled non-equilibrium condition. In addition, various structures in biomatrices across different scales will interfere with optical detection, especially for single molecule analysis.

Fluorescence correlation spectroscopy (FCS) and fluorescence recovery after photobleaching (FRAP) have been applied to investigate biomolecular diffusion in aqueous solutions as well as in 3D matrices. Different from aqueous solutions, the biomatrices provide very complex environments for the fluorescently labelled analytes, presenting diverse chemical groups of high concentrations and structures at different hierarchical levels. Consequently, FCS-based diffusion studies in the presence of biomatrices or cells will suffer from a low signal-to-noise ratio and significant batch-to-batch variation^8^. The photo-bleaching within the FRAP measurements that is used to induce non-equilibrium conditions is a very harsh process. The photochemical reactions can be greatly affected by the chemical surrounding associated with the biomatrices. We have observed that both photolysis and photo-bleaching can be drastically affected in different biopolymer systems. (supporting figure 1) Although physically more complex and less understood than normal diffusion, thermophoresis measurements can create non-equilibrium conditions without harsh photochemical reactions. We have found that the fluorescence measurement over a large area (∼ 10,000 *µ*m^2^) in the MST assay is tolerant to the optical features of various biomatrices used in this work.

**Figure 1.**
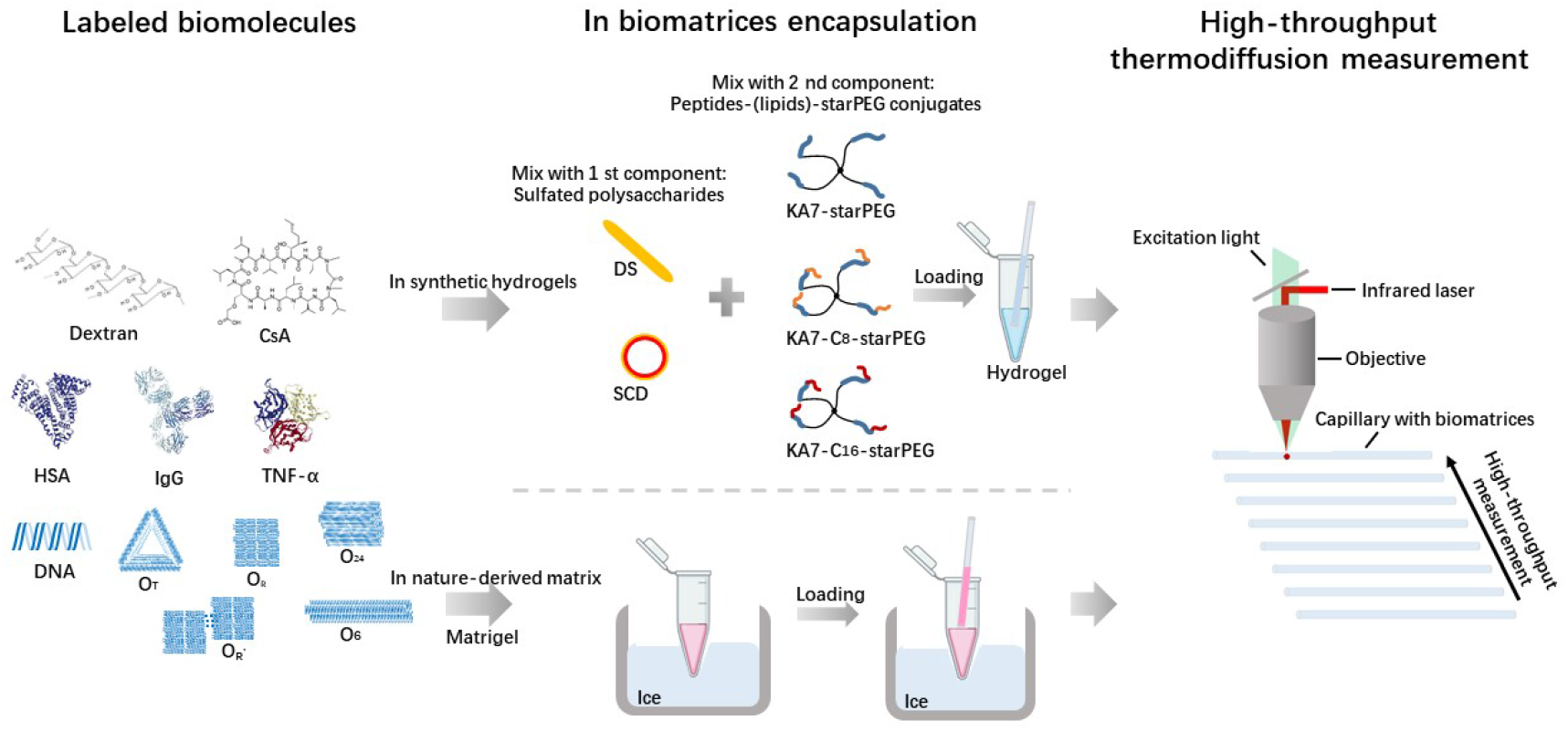
Preparation of biomolecules encapsulated in biomatrices for microscale thermophoresis (MST) measurements. First, biomolecules including polysaccharide (dextran), a small-molecule drug (CsA), proteins (HSA, IgG, TNF-α), DNA, DNA origami (O_T_, O_R_, O_R’_, O_6_, O_24_) are fluorescently labeled. They are further homogenously encapsulated into either synthetic hydrogels or nature-derived matrix. The liquid-like mixtures can be loaded into capillaries. After incubation, the hydrogel samples can be used in MST measurements. CsA, cyclosporine A; HSA, human serum albumin; IgG, immunoglobulin G; TNF-α, tumor necrosis factor α; O_T_, triangular DNA origami; O_R_, rectangular DNA origami without edge staples; O_R’_, rectangular origami with edge staples; O_6_, six-helix bundle DNA origami; O_24_, 24-helix bundle DNA origami; DS, dextran sulfate; SCD, sulfated cyclodextrin; KA7, seven repetitive lysine-alanine; starPEG, four-armed end-functionalized polyethylene glycol.

In this study, to apply MST assay to the study of different biomolecules in 3D matrices, we have employed two self-assembled hydrogel systems that allow for the homogeneous encapsulation of fluorescently labelled analytes without any chemical crosslinking reactions. Matrigel, a widely used ECM mixture derived from mouse tumor cells, represents a system compatible with such a study. It is a solution when kept on ice and can be mixed with desired reagents or cells. Upon incubation at elevated temperature (e.g. room temperature or 37 °C), a homogeneous gel can be formed. While Matrigel is a poorly defined protein mixture and has large batch-to-batch variation, a modular synthetic system, in which each component can be individually tuned, will allow for a more insightful understanding of analyte-matrix interactions. We have established a non-covalent hydrogel system formed by mixing heparin and peptide-starPEG (four-armed polyethylene-glycol) conjugate. The peptide has a minimal motif of (BA)_n_ (B is a basic residue, i.e., lysine (K) or arginine (R), A is alanine, and n is the number of repetitions). While the (BA)_n_ motif and a sulfated polymer represent the two prerequisites for hydrogel formation, the system is otherwise very flexible and tolerates different types of modifications, including extending the peptide sequences or using various types of the sulfated polysaccharide^9,10^. Recently, a lipophilic module has been incorporated into the system, resulting in a zwitterionic network containing amphiphilic structures. The synthetic system can mimic many chemical features of ECM by presenting polypeptide, oligosaccharide, and lipophilic moieties. It is important to note that different from covalently crosslinked polymers, these self-assembled matrices possess heterogeneous and large pores (in *µ*m scale)^11^, thus allowing for the diffusion of molecules and nanostructures. Owing to the high-throughput of the MST assay, we investigated the thermophoretic behaviors of a large variety of biomolecules of different natures and sizes (e.g. small molecules, polysaccharides, DNAs, DNA origami, proteins) in biomatrices of various compositions. Interestingly, we found that both the fluorescently labelled analytes and matrices can drift along temperature gradients. The thermodiffusion measurements in self-assembled biomatrices can give us insights into biomolecule-matrix interaction through a previously undescribed mechanism.

## Results and discussion

### Preparation of biomatrices and cell-laden hydrogel samples for MST measurements

To prepare peptide-starPEG/polysaccharide hydrogels for MST measurements (figure **1**), a fluorescence analyte was added to the polysaccharide solution, then mixed with the peptide-starPEG solution. The resulting mixtures remain fluidic shortly after mixing, allowing for adequate mixing and loading into glass capillaries. To prepare the Matrigel samples, fluorescently labelled analyte was added to Matrigel solution on ice (figure **1**). The mixture was then loaded into glass capillaries. The gel-loaded capillaries were incubated for 2 hours in a humidified chamber at room temperature for gelation, before subjecting to MST measurements (figure **1**). The procedures are also compatible with preparing cell-laden materials in the capillaries.

### Thermodiffusion of dextran in hydrogels

We first performed MST measurements with Cyanine(Cy5)-labelled 10 kD dextran (20 nM) in KA7-starPEG/dextran sulfate(DS) hydrogel (0.3 mM KA7-starPEG and 0.3 mM DS) and compared it with the measurements in aqueous solution (figure **2**). As thermodiffusion is balanced by ordinary diffusion in static state, constant diffusion and thermodiffusion coefficients both lead to an exponential depletion law and the Soret coefficient is defined as ratio ST = DT/D. The curves can be fitted to first-order reaction kinetics and the rate K and amplitude of relative fluorescence change (A_T_) can be calculated (supporting figure 2). While the K value in aqueous solution (0.1463 ± 0.0008) is higher than that in the hydrogel (0.0988 ± 0.0008), surprisingly, the A_T_ value in the hydrogel (0.2853 ± 0.0007) was found to be remarkably larger than that in aqueous solution (0.2205 ± 0.0006). Increasing the concentrations of KA7-starPEG and DS from 0.3 mM to 3 mM will cause a dramatic increase of storage modulus from 100 Pa to 100 kPa, as measured using a rheometer, while reduced K values and enhanced A_T_ values have been measured in the stiff hydrogel, as compared to the values in solution (figure **2B** and **2D**). Similar results of enhanced A_T_ values in the self-assembled network were obtained with the KA7-starPEG/sulfated cyclodextrin (SCD) hydrogel, KA7-C_16_-starPEG/DS hydrogel and KA7-C_16_-starPEG/SCD hydrogel. This observation is unexpected, as a matrix network should hinder the molecular motion. One explanation is that either KA7-starPEG or DS can interact with the analyte, thus changing its thermophoretic behavior. We performed the MST measurements of Cy5-dextran in either 0.3 mM KA7-C_16_-starPEG or 0.3 mM DS solution. Interestingly, no remarkable differences have been observed among these measurements in different aqueous solutions (figure **2B**). We speculated that the non-covalently assembled network might also undergo thermodiffusion, which could have an impact on the thermodiffusion of the fluorescence analyte (figure **2D**).

**Figure 2.**
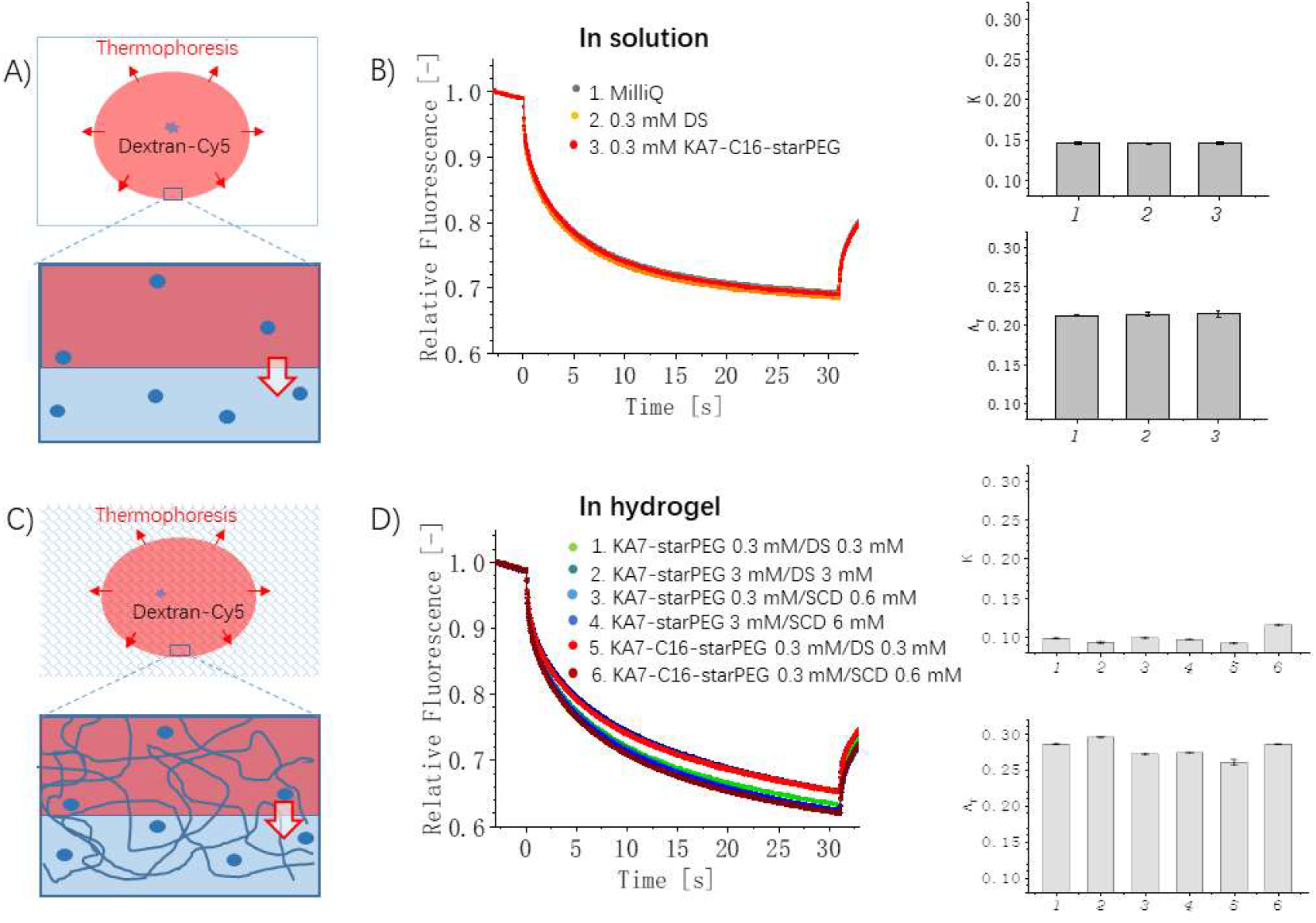
Thermodiffusion of dextran in aqueous media and matrices. A-B) The thermodiffusion of dextran in solution is represented by MST curves. The analysis is performed using two parameters, K and A_T_. C-D) Thermodiffusion of dextran in matrices is represented by MST curves, which are analyzed using parameters K and A_T_. K, rate; A_T_, amplitude of relative fluorescence change; DS, dextran sulfate; SCD, sulfated cyclodextrin; KA7, seven repetitive lysine-alanine; starPEG, four-armed end-functionalized polyethylene glycol. Error bars show the standard deviation (n=3).

### Thermodiffusion of hydrogel networks

To investigate whether the matrices can also drift along a temperature gradient, we performed MST measurements with KA7-starPEG/DS hydrogels doped with either Cy5-labelled KA7-starPEG or Cy5-labelled heparin (figure **3B** and figure **3C**). Like DS, heparin is a highly sulfated polysaccharide and can form stable hydrogel with KA7-starPEG. Indeed, the peptide-starPEG/sulfated polysaccharide hydrogel system was first developed for heparin^9^. As shown in figure **3B** and **3C**, both labelled KA7-starPEG and labelled heparin undergo thermodiffusion in the hydrogel, reflecting the motion of the entire network. The rate value K of KA7-starPEG and heparin are similar to each other. The amplitude value A_T_ of labelled KA7-starPEG is smaller than that of heparin, reflecting the lower mobility of the branched structure in a network. Moreover, the K and A_T_ values of labelled KA7-starPEG in KA7-starPEG solution are smaller than the values in the hydrogel, while the K and A_T_ values of labelled heparin in solution are also smaller than the values in hydrogel. Therefore, the formation of a non-covalently assembled network among the macromers does hinder their mobility. Because Matrigel is a heterogeneous mixture of myriads of different biomolecules, its network thermophoresis cannot be easily measured because of the lack of a simple labelling method. Nevertheless, the thermodiffusion of various biomolecules in Matrigel has exhibited a similar trend as in the synthetic matrices (as shown below).

**Figure 3.**
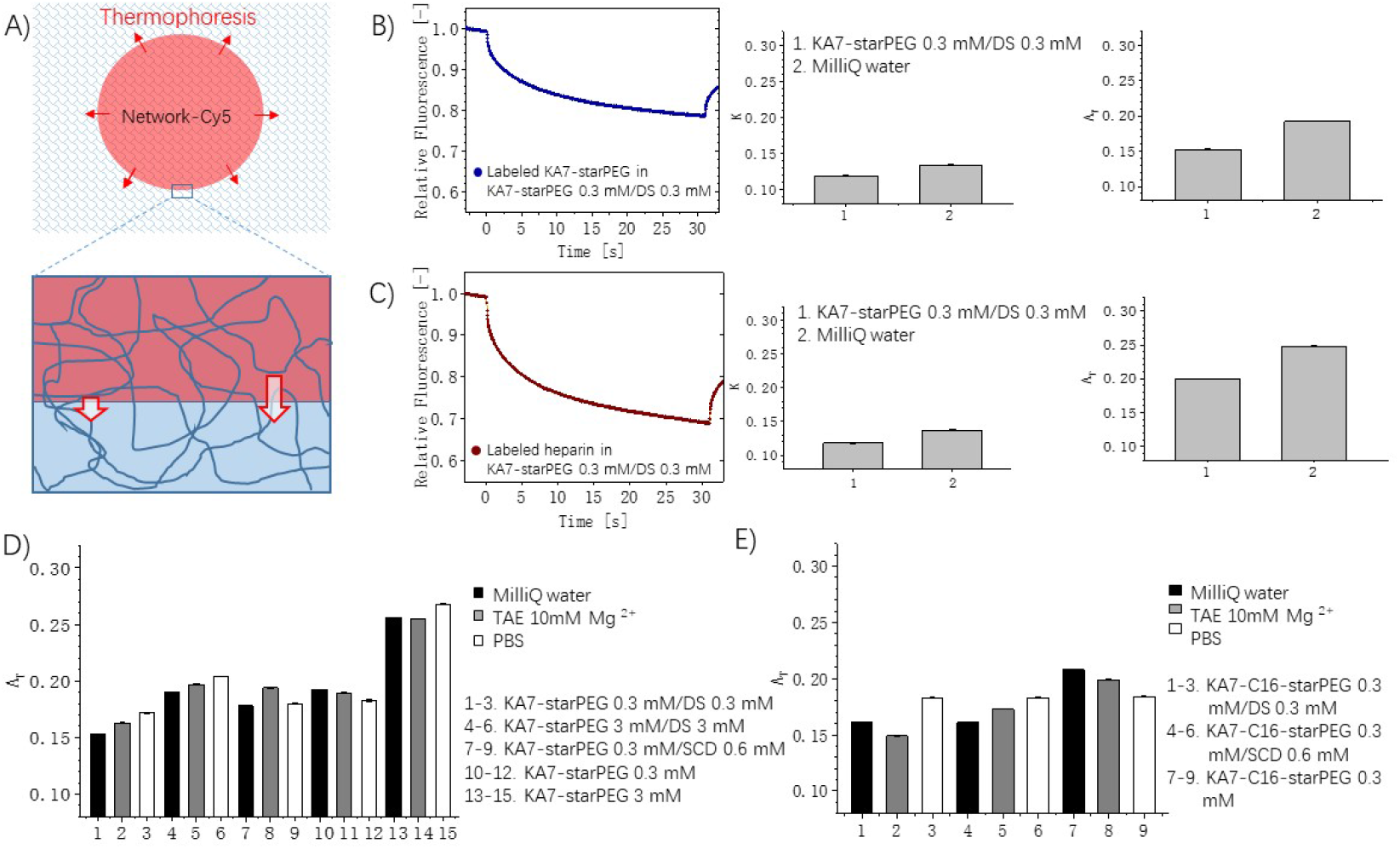
Thermodiffusion of hydrogel networks. A) Schematic represents thermodiffusion of hydrogel networks. The network components are fluorescently labelled. B) KA7-starPEG and C) heparin are fluorescently labelled and characterized by MST curves. The analysis of curves is performed using parameters K and A_T_. D)-E) Thermodiffusion of networks with labeled KA7-starPEG. DS, dextran sulfate; SCD, sulfated cyclodextrin; KA7, seven repetitive lysine-alanine; starPEG, four-armed end-functionalized polyethylene glycol; PBS, phosphate-buffered saline; TAE, tris-acetate-ethylenediaminetetraacetic acid. Error bars show the standard deviation (n=3).

Since the A_T_ value of labelled KA7-starPEG in matrix reflects the response of the network to a thermogradient, we next measured the network thermophoresis of KA7-starPEG/DS hydrogels with varied solid content. Interestingly, the A_T_ value of labelled KA7-starPEG in 3 mM hydrogel is higher than that in 0.3 mM hydrogel, but lower than its A_T_ value measured in 3 mM KA7-starPEG solution. Moreover, similar changes have been observed in water, in PBS buffer, and in TAE buffer (figure **3D**). While the buffer conditions can have a profound influence on thermodiffusion^5^, the network thermophoresis and the reduced A_T_ values in the network as compared to those in aqueous solutions have also been observed in different buffer systems. When DS is changed to sulfated cyclodextrin (SCD), matrix thermophoresis can also be observed. The effects of solid content and buffer condition on matrix thermophoresis were very small (figure **3D**). Matrix thermophoresis and reduced A_T_ values of labelled KA7-starPEG in a matrix as compared to that in solution have also been observed in KA7-C_16_-starPEG/DS and KA7-C_16_-starPEG/SCD hydrogels (figure **3E**).

The gelation of KA7-starPEG/DS and KA7-starPEG/heparin hydrogels is largely driven by the electrostatic interaction between the positively charged peptides and the negatively charge polysaccharides. While the KA7-lipid-starPEG/DS system introduces a lipophilic component into the zwitterionic matrix, the KA7-starPEG/SCD hydrogel possesses a relatively hydrophobic interior in the network^11^. Matrix thermophoresis has been detected in these materials of different compositions and solid contents. Moreover, when Cy5-dextran, a neutral hydrophilic macromolecule, was used as the analyte, remarkably enhanced A_T_ value has been measured in all these hydrogels of different compositions and solid contents (figure **2**), as compared to the A_T_ values in solutions. To investigate whether the additive effect of matrix thermophoresis on the thermophoresis of soluble analyte (figure **2D**) occurs also to other molecules, we performed MST measurements with other molecules (small molecules, polysaccharides, DNAs, DNA origami, proteins) in various hydrogels.

### Thermodiffusion of cyclosporin A in biomatrices

The natural product cyclosporin A (CsA) is a hydrophobic cyclic undecapeptide. In the design of drug formulations and delivery of this immunosuppressive drug, modulating its solubility is critical to achieve an optimal therapeutic effect^12^. Like dextran, the A_T_ value of Cy5-lablled CsA in 0.3 mM KA7-starPEG/DS hydrogel is larger than that of the matrix and that of the analyte in solution (figure **4A**).

**Figure 4.**
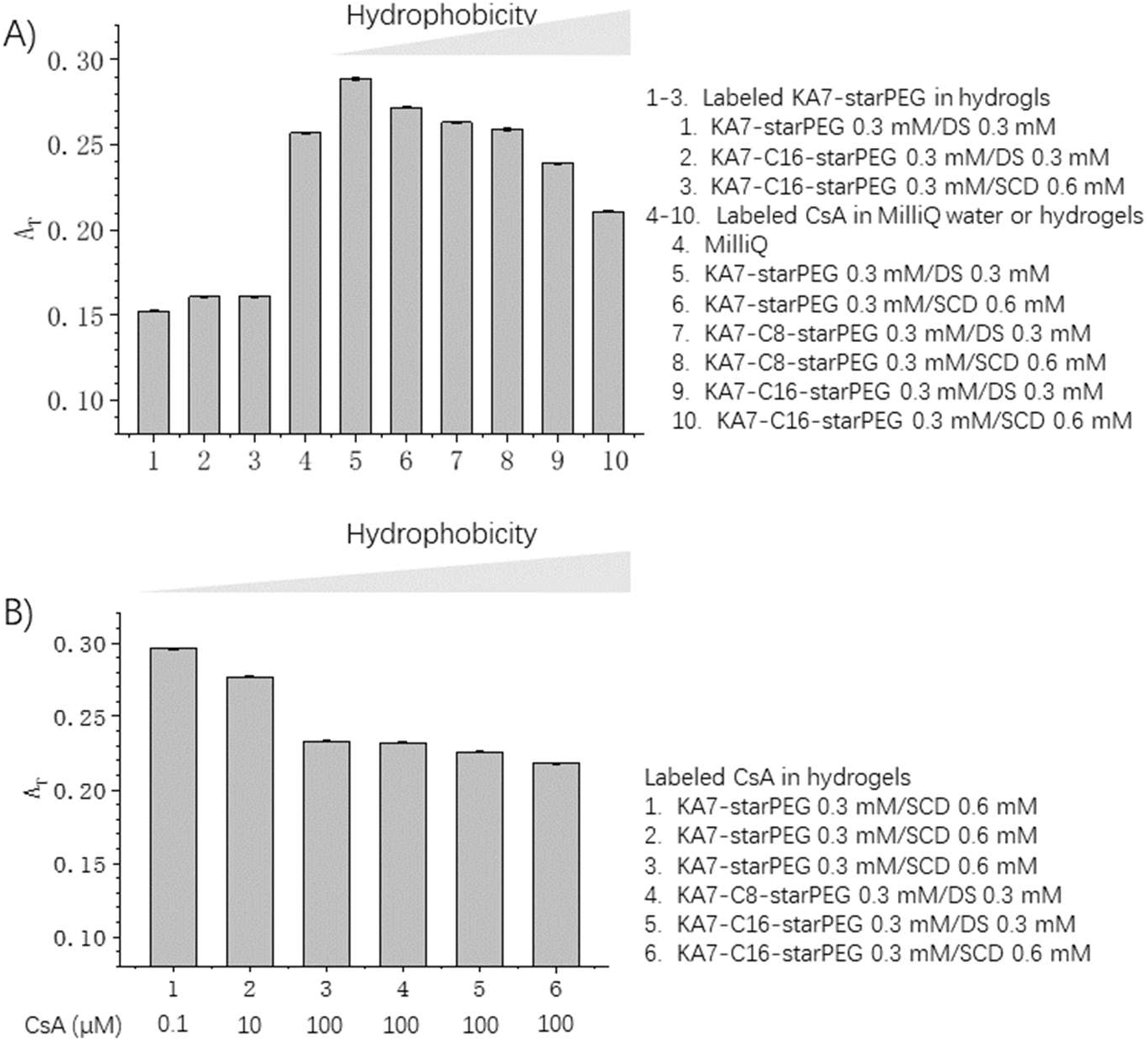
Thermodiffusion of CsA in hydrogels. A) Thermodiffusion of CsA in synthetic hydrogels with various hydrophobicity. B) Thermodiffusion of CsA in synthetic hydrogels with various hydrophobicity affected by both network components and the hydrophobic compounds themselves. CsA, cyclosporine A; DS, dextran sulfate; SCD, sulfated cyclodextrin; KA7, seven repetitive lysine-alanine; starPEG, four-armed end-functionalized polyethylene glycol. Error bars show the standard deviation (n=3).

We then investigated whether introducing a lipophilic group into the matrix can affect the A_T_ value of Cy5-CsA. Interestingly, with the increase of lipophilicity, a gradual decrease of A_T_ values has been measured (figure **4A**). Although the introduction of lipid group into matrix has little effect on the thermodiffusion of the network, the lipid moieties can interact with the hydrophobic compound, thus limiting its thermodiffusion as a free molecule. We have previously shown that Cy5-CsA can bind to KA7-C_16_-starPEG and KA7-C_8_-starPEG with *K*_d_ values of 30 *µ*M and 608 *µ*M, respectively^11^. Eventually, in an amphiphilic matrix, the A_T_ value of Cy5-CsA can be smaller than that in solution, especially in the KA7-C_16_-starPEG/DS and KA7-C_16_-starPEG/SCD hydrogels. Different from dextran that does not interact directly with both components in the self-assembled network, hydrophobic interactions with the matrix limit the thermodiffusion of Cy5-CsA upon encapsulation into an amphiphilic environment. Interestingly, while the *K*_d_ value between Cy5-CsA and KA7-starPEG is larger than 1 mM, reduced A_T_ value of Cy5-CsA (0.2596 ± 0.0006) has been observed in the 3 mM KA7-starPEG/DS hydrogel, as compared to that in 0.3 mM hydrogel (0.2888 ± 0.0006). The high A_T_ value of Cy5-CsA has been measured in Matrigel (0.3062 ± 0.0006), remarkably larger than that in solution (0.2570 ± 0.0007). However, as we cannot quantify the matrix thermophoresis of Matrigel, its effect on Cy5-CsA thermophoresis remains to be investigated in the future.

To mimic the condition of drug delivery with CsA-loaded biomaterials^13^, we performed MST measurements of Cy5-CsA with increasing concentration of non-labelled CsA encapsulated in the hydrogels. As the solubility of CsA in the aqueous buffer is around 10 *µ*M (figure **4B**), the thermodiffusion of Cy5-CsA was slightly affected with up to 10 *µ*M of CsA. Interestingly, at a concentration of 100 *µ*M CsA, the hydrophobic molecule can create an amphiphilic environment in the matrix, thus leading to a remarkable reduction of A_T_ value. Further increasing the hydrophobicity by introducing lipid chains and SCD to the hydrogel can cause a further reduction of the A_T_ values.

### Thermodiffusion of DNA and DNA origami in biomatrices

We then investigated the thermophoresis of Cy5-labelled double-stranded DNA and various DNA origami shapes. As expected, both KA7-starPEG/DS and KA7-starPEG/SCD hydrogels, as well as the hydrogels of different solid contents have caused increases in the A_T_ value of fluorescent DNA (figure **5A**). However, the effects are relatively small, as compared to the effect on Cy5-labelled dextran.

**Figure 5.**
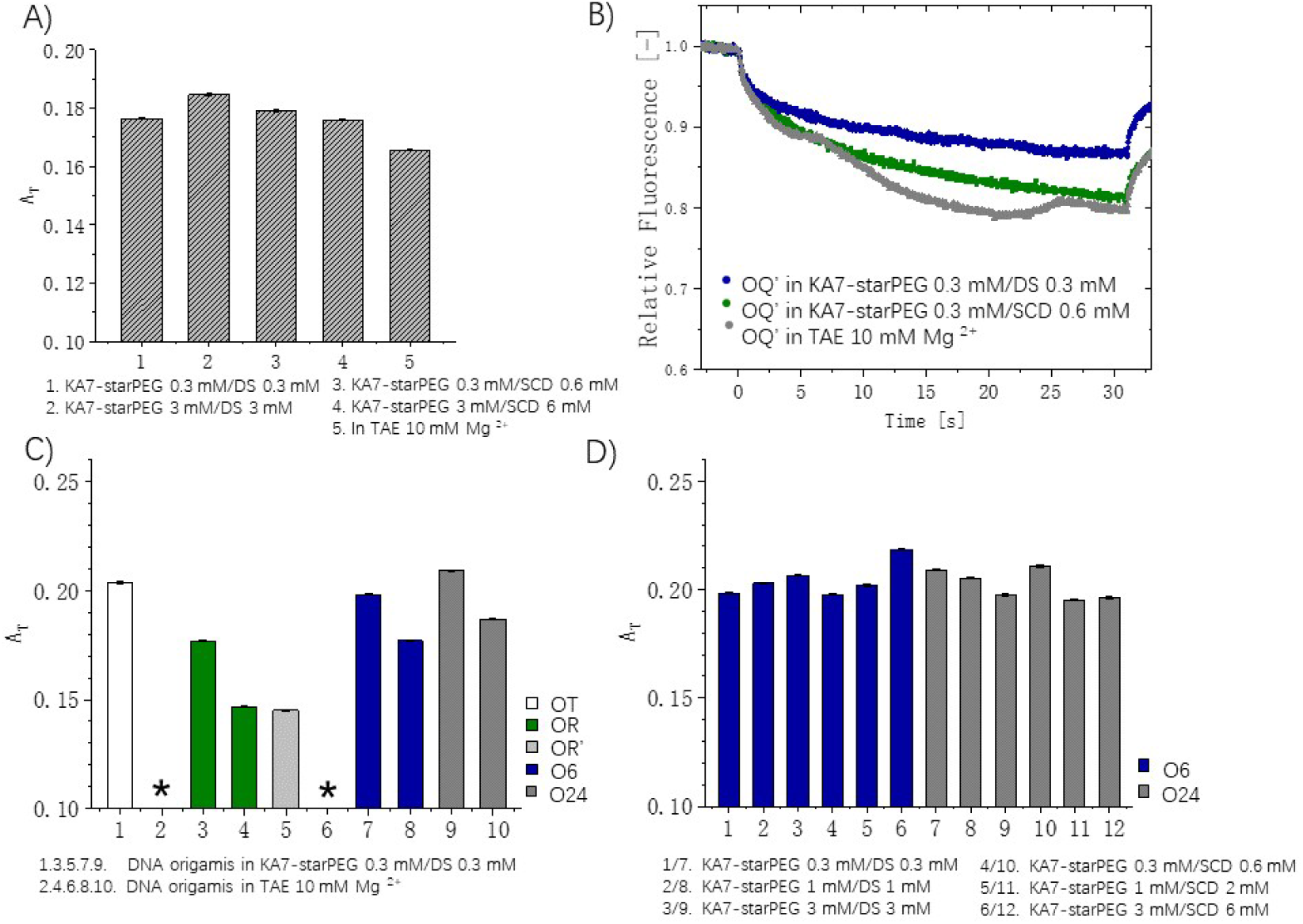
Thermodiffusion of DNA and DNA origami in solution and matrices. A) Thermodiffusion of DNA in solution and matrices. B) Aggregation of rectangular DNA origami O_Q_’ with edge staples can be prevented in the hydrogel network but not in solution. O_T_, triangular DNA origami; O_R_, rectangular DNA origami without edge staples; O_R’_, rectangular DNA origami with edge staples; O_6_, six-helix bundle DNA origami; O_24_, 24-helix bundle DNA origami; DS, dextran sulfate; SCD, sulfated cyclodextrin; KA7, seven repetitive lysine-alanine; starPEG, four-armed end-functionalized polyethylene glycol; TAE, tris-acetate-ethylenediaminetetraacetic acid. *Origami aggregation. Error bars show the standard deviation (n=3).

We have chosen five different DNA origami shapes for the MST study (figure **5B-D**). The triangular DNA origami O_T_ represents one of the most studied DNA nanostructures. The rectangular DNA origami O_R_ exhibits DNA blunt ends at its long edges, which facilitates aggregation via blunt-end stacking.^14^ Omitting the edge staples from the full staple set results in unpaired scaffold loops at the long edges, which prevents the aggregation of the rectangular DNA origami O_R_.^15^ While O_T_, O_R_, and O_R’_ are planar structures, the six-helix bundle (O_6_) and 24-helix bundle (O_24_) DNA origami are 3D structures.^16,17^ As expected, O_R_, but not O_R’_, forms aggregates in aqueous solution, thus preventing the MST measurement of O_R_ in aqueous solution (figure **5B**). While blunt-end stacking is minimized in the O_T_ design, it can nevertheless result in small aggregates via intermolecular contacts at the vertices, which interferes with the MST measurement in solution. The thermophoresis of O_R’_, O_6_, and O_24_ can be measured, and the curves can be fitted to first-order reaction kinetics (figure **5C**). Remarkably, upon embedding the nanostructures in hydrogels, the problem associated with aggregation can be remarkably suppressed (figure **5B**). As most biomolecular aggregation is mediated by weak intermolecular interactions, segregating them with a polymer matrix may diminish the probability of their interactions. The segregation effect of hydrogels (including Matrigel) has also been observed for other types of aggregating biomolecules (e.g., TNF-α, as shown later).

The A_T_ values of O_T_ and O_R_ in KA7-starPEG/DS hydrogel cannot be directly compared with those in aqueous solution because of aggregation. While the A_T_ values of O_R’_, O_6_, and O_24_ in aqueous solution differ from each other remarkably, a similar increase has been measured for all these structures upon encapsulation in KA7-starPEG/DS hydrogel (figure **5C**). Moreover, increasing the hydrogel solid contents has a moderate effect on the increased A_T_ values, as measured for O_6_ and O_24_ (figure **5D**). Interestingly, whereas the A_T_ values of O_6_ in hydrogels increase upon increasing the solid contents, opposite changes have been measured for O_24_. It is important to note that the thermodiffusion of DNA origami in the hydrogels is larger than in solution as well as the thermodiffusion of hydrogel, reflecting the ability of the nanostructures to move through the porous network (figure **2D**).

### Thermodiffusion of proteins in biomatrices

Pro-inflammatory cytokine TNF-α, human serum albumin (HSA) and immunoglobin G (IgG) have been chosen for studying protein thermophoresis in self-assembled polymer matrices. While TNF-α represents one of the most valuable therapeutic targets^18,19^, HSA and IgG are the most abundant proteins in the bloodstream. Protein aggregation and adsorption on glass walls are often observed in MST measurements, which can be partially suppressed by using treated glass surfaces to prevent unspecific protein adsorption. Because of the segregation effect of hydrogels, the MST measurements can be reproducibly performed with various proteins, even when a protein such as TNF-α forms aggregates in the treated capillaries (Figure **6A**). As the proteins have little contact with the glass surface, noise associated with protein adsorption can be completely abolished even with untreated glass capillaries.

**Figure 6.**
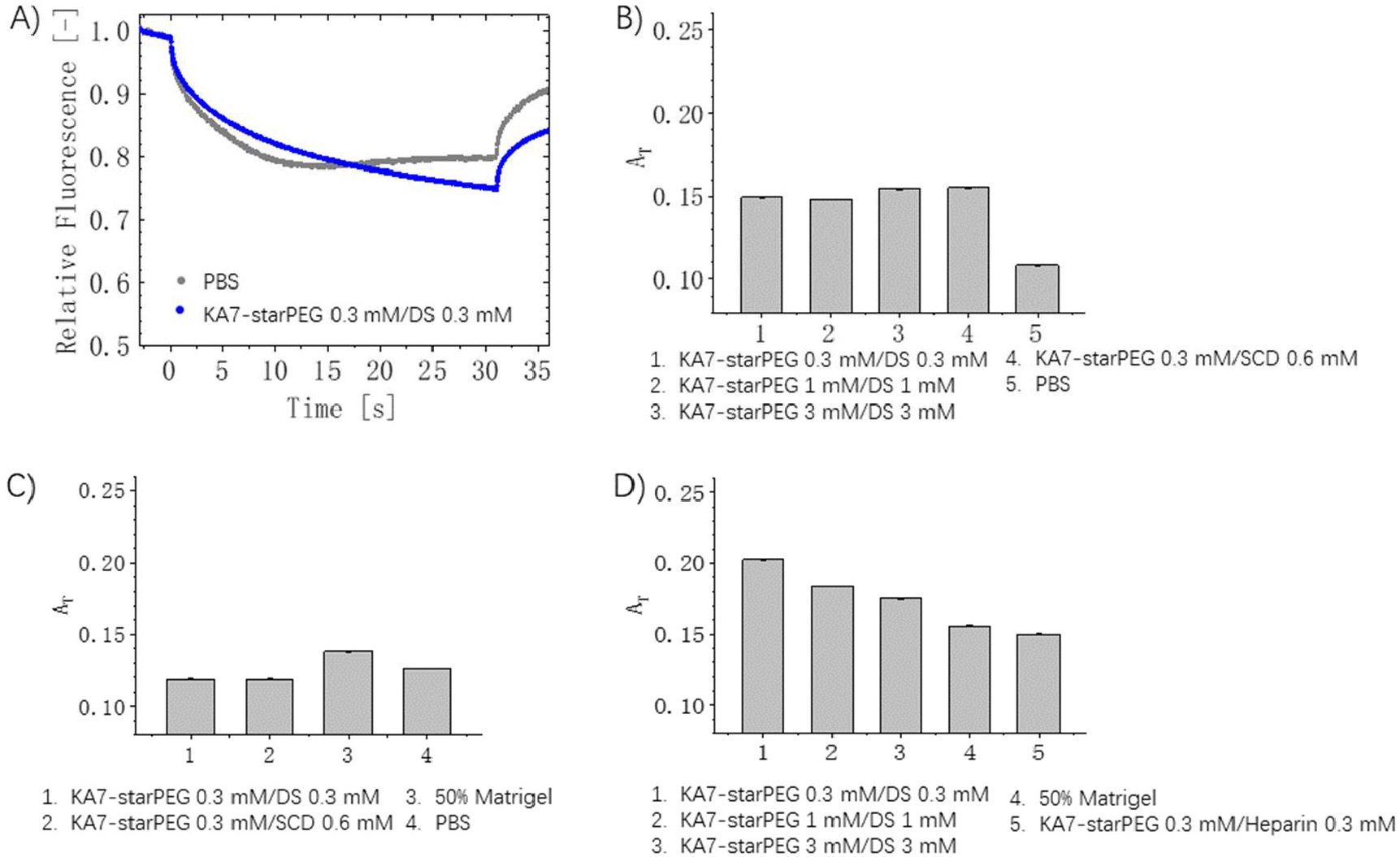
Thermodiffusion of proteins in solution and hydrogels. A) The aggregation of TNF-α (in buffer) can be prevented by the encapsulation in hydrogel. MST measurements of B) Albumin, C) IgG, D) TNF-α, in solution and biometrics. HSA, human serum albumin; IgG, immunoglobulin G; TNF-α, tumor necrosis factor α; DS, dextran sulfate; SCD, sulfated cyclodextrin; KA7, seven repetitive lysine-alanine; starPEG, four-armed end-functionalized polyethylene glycol; PBS, phosphate-buffered saline. Error bars show the standard deviation (n=3).

Like dextran and DNA origami, fluorescently labelled HSA has shown A_T_ values in hydrogels higher than the A_T_ value in solution (figure **6B**). Interestingly, while the A_T_ value of matrix thermophoresis (figure **3**) is larger than the A_T_ value of HSA in an aqueous solution, the remarkably increased A_T_ values of HSA do not exceed that of matrix thermophoresis. Encapsulation in hydrogel caused minor changes of the thermodiffusion of IgG (figure **6C**). While a small increase has been measured in Matrigel, small decreases have been measured in KA7-starPEG/DS hydrogel and KA7-starPEG/SCD hydrogel.

When TNF-α is encapsulated in hydrogels, all MST curves can be fitted to first-order reaction kinetics (figure **6D**). Different from HSA and IgG, the thermophoresis of TNF-α is greatly affected by hydrogel composition. Increasing the solid content of KA7-starPEG/DS hydrogel causes a reduction in A_T_ value. TNF-α possesses a heparin-binding domain. We have previously shown that the homotrimer of TNF-α can bind to and be stabilized by heparin.^20^ Interestingly, replacing DS with heparin caused a remarkable reduction of the A_T_ value in KA7-starPEG/heparin hydrogel. Both being able to bind to matrices through multivalent interaction, the thermodiffusion of fluorescently labelled KA7-starPEG in KA7-starPEG/DS hydrogel is very similar to that of fluorescently labelled trimeric TNF-α in KA7-starPEG/heparin hydrogel. The strong interaction between TNF-α with heparin makes the protein an integral part of the matrix. Interestingly, in Matrigel, the A_T_ value of TNF-α is higher than that in KA7-starPEG/heparin hydrogel, but remarkably lower than that in KA7-starPEG/DS hydrogel, indicating the presence of strong interaction between the cytokine with the native ECM material.

### MST measurements in the presence of cells

We then studied whether the MST assay of drug molecules such as CsA in hydrogels can be performed in the presence of cells. The AT value in cell culture media is remarkably lower than that in simple buffer systems, as CsA can interact with many different molecules in media, resulting in altered thermophoretic behaviors. Hydrogels with fluorescently labelled CsA were formed with cell culture media (figure 7A). Similar to the hydrogels formed in simple buffers, encapsulation of CsA in 0.3 mM KA7-starPEG/DS hydrogel in cell culture media resulted in a remarkable increase of their AT values (figure 7B).

**Figure 7.**
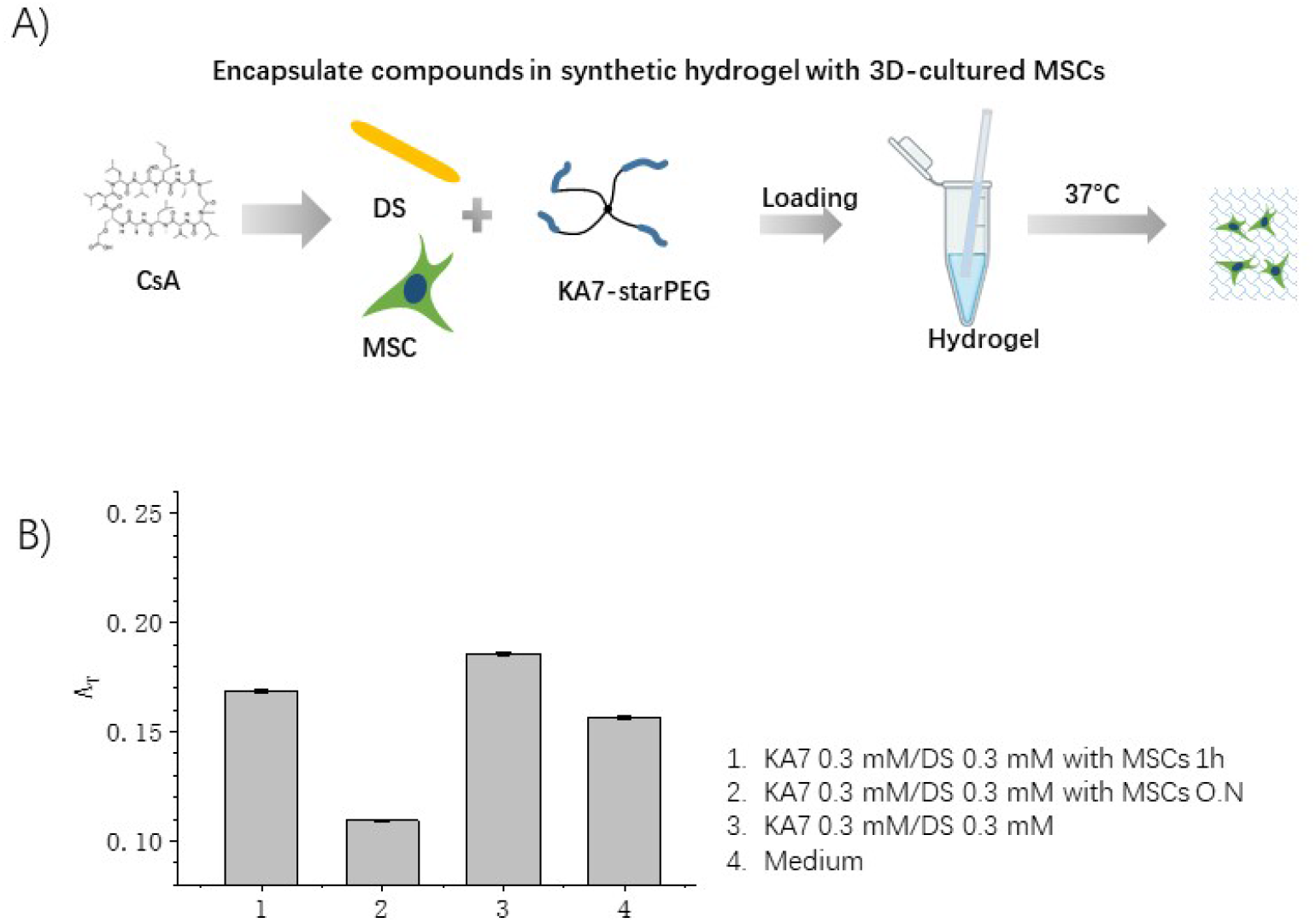
Thermodiffusion of CsA in the synthetic hydrogels with 3D-cultured MSCs. A) Preparation of biomolecule-embedded biometrics in the presence of MSCs for MST measurement. B) MST measurements of fluorescently labelled CsA in cell culture media, in hydrogel, and in cell-laden hydrogel. CsA, cyclosporine A; DS, dextran sulfate; MSC, mesenchymal stromal cell; KA7, seven repetitive lysine-alanine; starPEG, four-armed end-functionalized polyethylene glycol; h, hour; O.N., overnight. Error bars show the standard deviation (n=3).

We then formed the hydrogels in the presence of MSC to perform the MST measurements (figure 7A). Interestingly, while all curves can be fitted to first-order reaction kinetics, the presence of cells has caused remarkably decreased AT values, though the cells only contribute to a small fraction of the entire hydrogel mass (cell number 10^5^ cells/ml). The drug molecule can interact with the cells, thus affecting their thermophoretic behaviors. Interestingly, after culturing the cell-laden materials in capillary overnight, a further remarkable decrease in AT value has been measured (figure 7B). CsA can enter into cells, resulting in confinement to limit its movement. It is important to note that the cells are alive under this condition. Although it is difficult to fully characterize the cell viability in capillaries, their morphology and migration to form clusters resembled the observations in our previous studies.^21^

## Conclusion

We have demonstrated that MST assays can be performed with either natural or synthetic hydrogels, showing high reproducibility, and the curves can be fitted to first-order reaction. Various biomolecules with hundred-fold variation in their sizes and million-fold variation in their volumes, can be analyzed in various matrices. Remarkably, the presence of cells does not interfere with the measurements, whereas their interaction with fluorescently labelled biomolecules affects the thermophoretic behavior of analytes. While thermodiffusion has been known for a long time^2^, the underlying physics remains poorly understood^3^. Nevertheless, the phenomenon has been successfully used as an all-optical technique to measure molecular interactions^22^. In this work, we have expanded the assay to 3D matrices, which are more biologically relevant than aqueous solutions, to resemble the native environments of biomolecules in tissues. Interestingly, we found that the matrices can also drift along the temperature gradient, thus enhancing the thermophoresis of many analytes. The thermal gradient of a few degrees generated by an IR laser in MST measurements represents a very mild method to create a non-equilibrium condition, not only of the fluorescently labelled analyte, but also of the surrounding network. Further study to understand the physics of molecule and matrix thermophoresis could help us to characterize some fundamental biochemical processes including binding and diffusion in various complex biological systems with a simple MST measurement.

While proteins are structurally more diverse and complex than other classes of biomolecules, their aggregation represents a common problem for many biochemical assays. We have shown that the segregation of molecules by polymer networks can diminish the formation of aggregates in the MST assays. The use of MST to study protein-biomatrix interaction remains to be explored in the future. While the thermodiffusion of HSA is little affected by matrix composition, the hydrogel composition exhibits a remarkable effect on the A_T_ values of TNF-α. IgG is the only molecule in this study, whose thermodiffusion is not enhanced upon encapsulation into hydrogels. In addition to the nature of biomolecules and the aqueous environment, their interaction with matrix, as well as the back-flow of water molecules to compensate for the expanding network can all affect the measured values.

## Acknowledgments

We thank Ulrike Hofmann for technical support. This project was supported by the German Bundesministerium für Bildung und Forschung (BMBF) Grant 03Z22E511.

