## Supplementary material for "Thermophoresis of Molecules and Structures of different Sizes in Self-assembled Biomatrices": Methods and supporting figures

### Materials and methods

1. Assembly and purification of peptide-(lipid)-starPEG.
All the assembly and purification methods of peptide-polymer conjugates are published in doi:10.1021/acs.chemmater.0c04105.

3. Assembly of DNA origamis and atomic force microscopy (AFM).
All designs are published as standard assembly conditions. Assembly conditions for the triangles, rectangles, and 6HBs are the same as in doi:10.3390/nano10112200. The 24HBs were assembled according to the protocol described in doi: 10.1093/nar/gkab097.

2. DNA double-strand P2 pair was purchased from Sigma-Aldrich.

5’-GAGATCGGAAGAGCGTCG-3’
3’-CTCTAGCCTTCTCGCAGC-5’

4. Diffusion of biomatrices or biomolecules in hydrogel measurement by MST
Cy5 labeled network compounds or biomolecules (or the mixture of CsA-Cy5/CsA) were prepared in DS or β-CD sulfate solutions (mixture 1). After mixing peptide(-lipid)-polymer conjugates solution with mixture 1, the mixtures were immediately loaded onto the MST premium standard glass capillaries. After overnight incubation in a high humidity environment (R.T.), the capillaries were measured with MST power of 20% and LED power = 40%. Each sample was repeated in triplicate and the concentration of the components was adjusted as described in the result section.

5. Diffusion of biomolecules in cell loaded hydrogel measurement by MST
Cy5 labeled CsA were prepared in DS solutions (mixture 1). Mesenchymal stromal cells were loaded in peptide(-lipid)-polymer conjugates solution (mixture 2). After mixing mixture 1 with mixture 2, the liquid-like mixture was immediately loaded onto the MST premium standard glass capillaries. After overnight incubation in a high humidity environment (R.T.), the capillaries were measured with MST power of 20% and LED power = 40%. Each sample was repeated in triplicate and the concentration of the components was adjusted as described in the result section.

**Supporting figures**

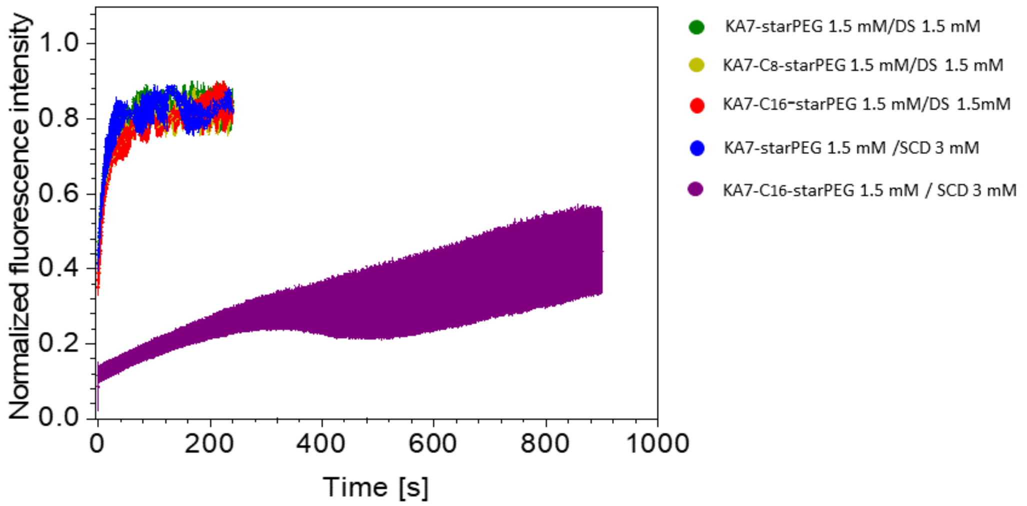


**Supporting figure 1.** Diffusion of CsA-Cy5 in biometrics measured by FRAP. FRAP measurements in synthetic hydrogels are very noisy, and the photo-bleaching reaction can be unexpectedly affected by matrix compositions.

In a previous study, we have synthesized a peptide containing photo-cleavable linker (PL) CWGG-PL-KA7. The peptide can by quickly cleaved upon exposure to light of 366 nm (5 min, with a hand UV lamp). The photolysis was followed by HPLC. The peptide CWGG-PL-KA7 was conjugated to maleimide starPEG, resulting in CWGG-PL-KA7-starPEG. The peptide-polymer conjugate can also be quickly cleaved upon exposure to light of 366 nm. However, upon forming hydrogels using CWGG-PL-KA7-starPEG with either heparin or dextran sulfate, the resulting hydrogels cannot be destroyed by long exposure to UV light (for more than 2 hours). Like the photo-bleaching reaction, the photolysis reaction can also be remarkably affected in a hydrogel environment. The underlying chemistry remains to be investigated in the future.

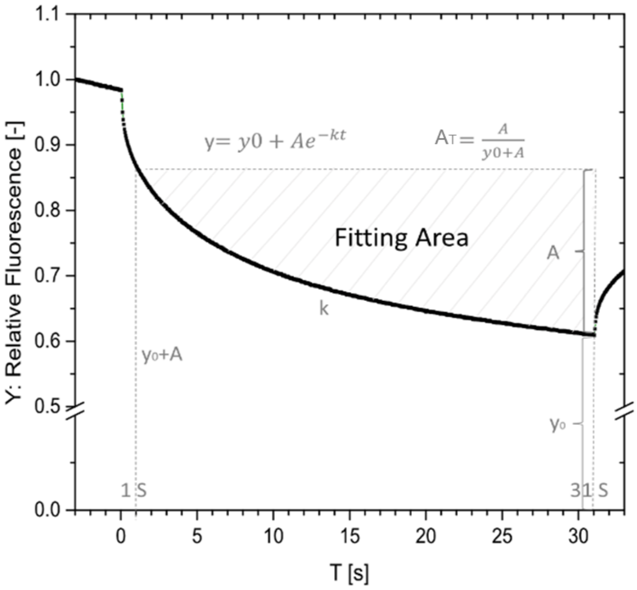


**Supporting figure 2.** The MST curves are fitted to first-to-order reaction kinetics and the rate K and amplitude of relative fluorescence change ($A_T$) can be calculated.

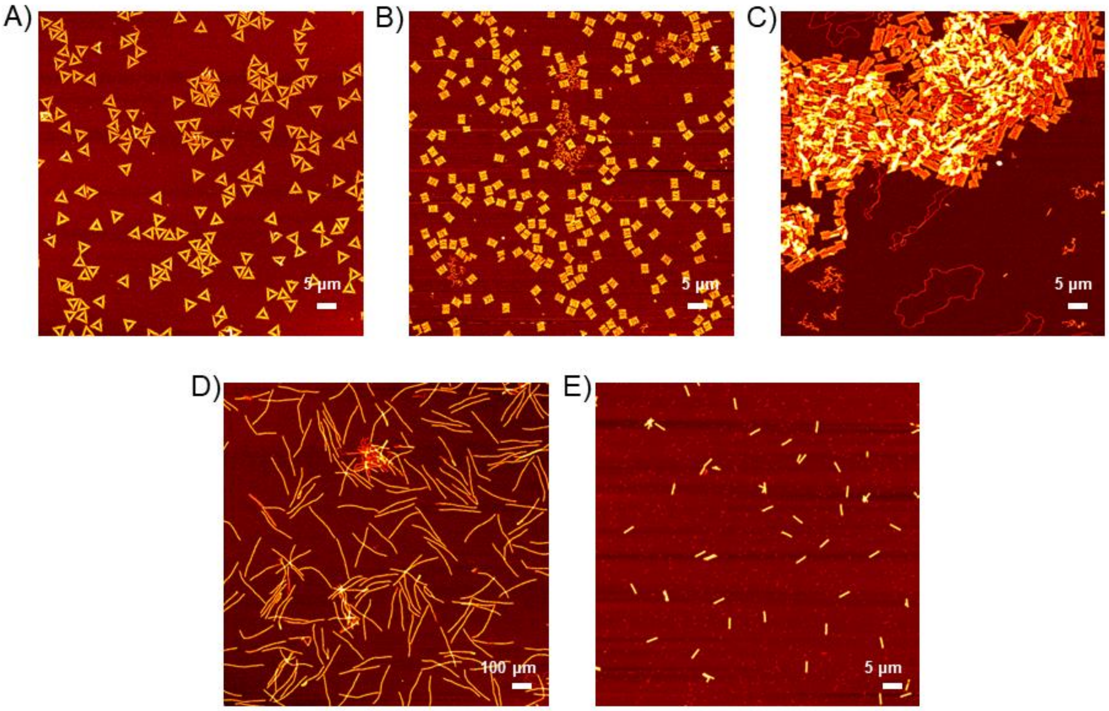


**Supporting figure 3**. The atomic force microscopy (AFM) images of the DNA origami. A) Triangle DNA origami, B) rectangle DNA origami, C) aggregated rectangle DNA origami, D) 6-helix bundle DNA origami, E) 24-helix DNA origami.
